## Supplementary material for "Anxious vision: Trait-like visual cortical hyperactivity in trait anxiety": SOM

**This PDF file includes:**

Supplemental analyses

Fig. S1

Table S1

References

### Experiment 1

**Additional Analyses**

Correlation between VEPs and trait anxiety for CS+ and CS- stimuli

Effects of conditioning on VEPs were reported in our earlier study (*1*). Here, to evaluate the effect of conditioning on the correlation between trait anxiety and P1/C1-N1, we also examined VEPs for CS+ and CS- stimuli separately at all 3 time points. As shown in **Table S1**, correlation was comparable for the two types of stimuli at all 3 time points, ruling out the effect on conditioning.

**Table S1 correlation between VEPs and trait anxiety for CS+ and CS- stimuli**

| Correlation coefficients | | Time | | |
|---|---|---|---|---|
| | | Time 1 (baseline) | Time 2 (Day 1 post-conditioning) | Time 3 (Day 16 Post-conditioning) |
| M-selective stimuli (P1) | CS+ | -0.356* | -0.383* | -0.334* |
| | CS- | -0.356* | -0.383* | $-0.316^{+}$ |
| P-selective stimuli (C1-N1) | CS+ | 0.351* | 0.374* | 0.379* |
| | CS- | 0.370* | 0.384* | 0.358* |

**p<0.05, +p<0.1.*

Comparison of correlation between trait anxiety and ERP components at different times

The correlation coefficients between trait anxiety and P1/C1-N1 at 3 time points were compared using Hittner's method based on Monte Carlo simulation (*2*). The analysis revealed no significant differences in the correlation coefficients between trait anxiety and P1 components across the three time (*p*'s > .44). Similarly, another analysis indicated no significant differences in the correlation coefficients between trait anxiety and C1-N1 components across the three times (*p*'s > .60). Therefore, the strength of the correlations remained consistent over time.

Point-by-point analysis of correlation between VEPs and trait anxiety

To delineate the precise time course of the association between VEPs and trait anxiety, we correlated amplitude at each data point within the first 200 ms with trait anxiety. We applied a permutation package in R that was designed to overcome Type I error from multiple comparisons (*3*) to determine the minimal window of consecutive data points with significant correlation ($p < 0.05$). Accordingly, at Time 1 (pre-conditioning baseline), significant time windows for M- and P-selective stimuli were identified as 74-200 ms, $p = 0.005$ and 86-175 ms, $p = 0.011$, respectively. At Time 2 (immediately post-conditioning), significant time windows for M- and P-selective stimuli were identified 42-200 ms, p = 0.002 and 86-171 ms, $p = 0.009$, respectively. At Time 3 (Day 16; 15 days post-conditioning), significant time windows for M- and P-selective stimuli were identified as 78-200 ms, $p = 0.02$ and 86-160 ms, $p = 0.029$, respectively.

<u>Correlation between P1 and C1-N1</u>

The intriguing opposite patterns of correlation between trait anxiety and the P1 versus C1-N1 prompted us to explore the direct association between P1 and C1-N1. As shown in **Fig. S1**, a significant negative correlation was observed between P1 and (inverted) C1-N1 amplitudes at every time point, $r = -0.62$, $p < 0.001$, $r = -0.73$, $p < 0.001$, $r = -0.63$, $p < 0.001$ at Time 1, Time 2, and Time 3, respectively.

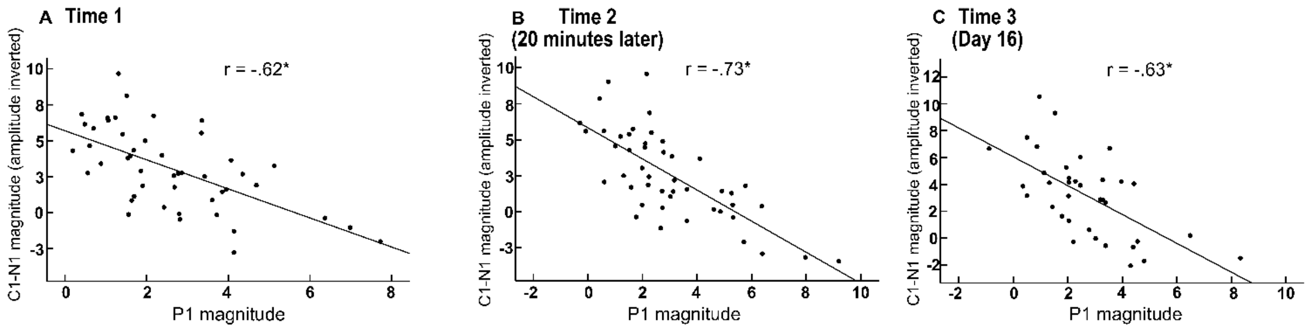


**Figure S1 Correlation between P1 and C1-N1 at each time point**

#### Experiments 2 & 3

##### Additional Analysis

<u>Point-by-point analysis of correlation between VEPs and trait anxiety</u>

Similar to **Experiment 1,** we conducted point-by-point correlational analyses between VEP potentials and trait anxiety in the first 200 ms, delineating the precise time course of this association. We applied a permutation package in R that was designed to overcome Type I error from multiple comparisons (*3*). In **Experiment 2**, a significant time window was detected for the P-selective stimuli (58-105 ms, p = 0.038), but not for the M-selective stimuli (**Figure 2B**; grey box at the bottom). Similarly, in **Experiment 3**, a significant time window was detected for the P-selective stimuli (46-94 ms, $p = 0.023$), but not for the M-selective stimuli (**Figure 2D**; grey box at the bottom).

<u>Comparison of correlation between trait anxiety and VEPs in **Experiments 2** and **3**</u>

We further compared the correlations between trait anxiety and P1/C1 in **Experiments 2** and **3**. We observed no difference in correlation coefficients between the two experiments, for either the correlation of trait anxiety with P1 amplitude ($p = 0.82$) or with C1 amplitude/magnitude ($p = 0.64$).

#### Experiment 4

##### Additional Analysis

<u>Point-by-point analysis of correlation between VEPs and trait anxiety</u>

Similar to the other experiments, we conducted point-by-point correlational analyses between VEP potentials and trait anxiety, using a permutation package in R to overcome Type I error from multiple comparisons (*3*). A significant time window was detected for the P-selective stimuli (101-140ms, $p = 0.047$), but not for the M-selective stimuli (**Figure 3B**; grey box at the bottom).

<u>Effects of emotion on correlation between VEPs and trait anxiety</u>

To explore whether the correlation between VEPs and trait anxiety was affected by emotion of the images, we conducted an analysis of covariance (ANCOVA) of emotion, SF, and trait anxiety on the VEPs. There was no interaction of emotion with SF and trait anxiety ($p = 0.22$). Specifically, the correlation between P1 and trait anxiety was comparable for fear, disgust, and neutral images in HSF: $r = 0.37$, 0.37, 0.36, respectively ($p$'s < 0.02), and similarly negligible for fear, disgust, and neutral images in LSF: $r = 0.21$, 0.16, 0.08, respectively ($p$'s > 0.20).
